# The immune map of human body

**DOI:** 10.1101/2024.03.03.583227

**Authors:** Rui Gan, Xiangxing Jin, Xianwen Ren

## Abstract

The immune system serves as a vital defense mechanism against diseases and infections for human bodies, comprised of a complex network of diverse cells, organs, and tissues. Here, we conduct single-cell deconvolution of 17382 RNA-seq samples from 30 tissues and construct a comprehensive map of the distributions of >1000 different cell states across all the major immune cell types throughout human bodies. This high-resolution immune map covers both sexes and a wide range of ages, enabling interrogation of the immune responses to sex differences and age-related changes. Similarity analysis among the different tissue types depicts the tissue-specific immunity of testis, liver, lung, brain, and kidney besides blood and spleen, and the high immune similarity between breast and adipose tissues for example. This comprehensive immune map may deepen our understanding of human immune system from a holistic perspective.

## Introduction

The immune system works in unison to safeguard the body against infections, acting as an integrated and highly developed network of diverse cell types to maintain tissue homeostasis and overall health ^1^. Each immune cell type plays a unique role in performing specific functions and communicating with other cells together to mediate protective immunity for the body. The major immune cell types are traditionally classified as lymphoid cells including B cells, T cells, natural killer (NK) cells as well as natural killer T (NKT) cells and myeloid cells including monocytes/macrophages (mono/macro), dendritic cells (DC), neutrophils, eosinophils, basophils, and mast cells ^2^. The lymphocytes mainly play pivotal roles in adaptive immunity, responsible for antibody production and direct killing of cells expressing specific antigens with immunologic memory ^3^. Meanwhile, myeloid cells play major roles in innate immunity, working in tandem to contain invading microorganisms and secrete different types of cytokines ^4^. In addition, endothelial cells are critical in immune regulation of limiting and facilitating the transport of various immune cell populations out of the vasculature and into tissue ^5^.

In recent years, single-cell RNA sequencing (scRNA-seq) has revolutionized the studies of human organs and tissues with accurate estimation and characterization of different cell types. Though previous large-scale studies have investigated the human immunity via scRNA-seq technologies, only one or a few tissues were studied due to the high-cost limitation of current technologies ^6,7^. Therefore, there is an urgent need for creating a comprehensive immune map of the human body across various immune cell types. Bulk RNA-seq is a straight-forward method for characterizing the averages of cellular expression, but fails to capture the cellular heterogeneity within samples ^8^. These challenges make it difficult to have a holistic understanding of human immunity across all tissues at cellular resolution ^6^.

Recently, multiple effective methods have been developed to reconstruct RNA-seq data by computational deconvolution with scRNA-seq data as reference such as Redeconve, an effective and efficient algorithm for estimating the cellular composition of spatial transcriptomics data at single-cell resolution ^9^. Inspired by the previous research efforts, we argue that integrative analysis of scRNA-seq data with bulk RNA-seq data could probably provide an effective solution to this problem. Here, we apply Redeconve on The Genotype-Tissue Expression (GTEx) to reconstruct the cellular compartments of 17382 bulk RNA-seq samples spanning 30 tissues ^10^, by using Tabula Sapiens ^7^ composed of 483,152 cells from 24 human tissues and organs as reference. By conducting such analysis, we successfully construct the most comprehensive immune landscape of the human body until now, providing a valuable data resource for the overall distribution of different immune cell types/states across most human tissues. This comprehensive immune map also enables investigating the associations of immune compositions with age and sex.

## Results

### Overview of the human immune map construction process

For convenience, we term this comprehensive human immune map as RedeImmunoMap, which is constructed by single-cell deconvolution of GTEx data with Redeconve as the algorithmic workforce and Tabula Sapiens as the single-cell reference. RedeImmunoMap presents a high-resolution immune map of human body by single-cell deconvolution of RNA-seq data (**Fig. 1**). Taking into account that Tabula Sapiens is a comprehensive and extensive human reference atlas as well as GTEx has generated a wealth of tissue samples from multiple donors with detailed information about age and sex, we integrated two datasets to reconstruct the cellular composition of immune plus endothelial cells for 17382 samples across 30 human tissues by single-cell deconvolution ^9^. Because the number of single cells is overwhelming, we totally took 5002 cell states sampled from 483,152 cells by originally annotated cell type ^7^. Thus, we obtained a high-resolution abundance matrix of 5002 cell states within 17382 samples, encompassing an extensive range of human tissues, including up to 2642 brain samples and at least 9 fallopian tube samples (**Fig. 1**). After picking immune and endothelial cells from 2920 detected cell states by their signature genes and combination with the original annotations, 999 immune and 248 endothelial cell states were totally detected and annotated as 12 major immune cell types including B cells, plasma cells, T cells, NK cells, NK T cells, monocytes/macrophages, DC, neutrophils, eosinophils, basophils, mast cells and innate lymphoid cells (ILCs) plus endothelial cells, subsequently clustered into 33 cell clusters (**Fig. 2A-C**). The averaged accuracy of the estimated abundance (defined as the cosine similarities between the true RNA-seq gene expression vectors and the reconstructed gene expression profiles) exceeded 0.9 for most tissues and reached 0.85 for all tissues (**Fig. 2D**), hopefully offering unprecedented insights into the intricate immune cellular composition of the human body.

**Fig. 1.**
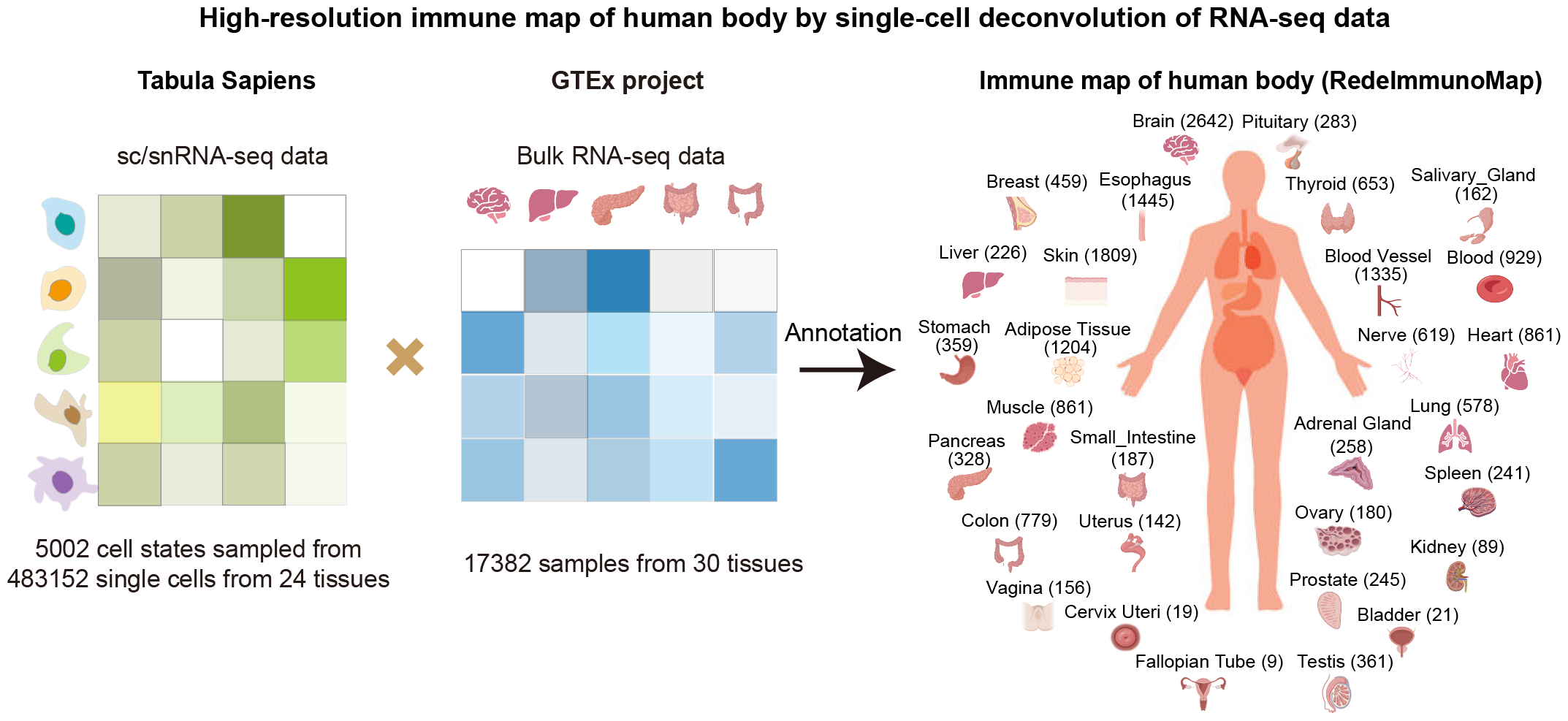
Overview of RedeImmunoMap. The scRNA-seq data from Tabula Sapiens and bulk RNA-seq data from GTEx were integrated by Redeconve. The estimated cell abundance matrix was analyzed to depict the immune abundance of human body across 30 tissues and organs. The largest population of tissue samples were brain samples and the smallest population of tissue samples were fallopian tube samples. The tissue figures were created with BioRender.com.

**Fig. 2.**
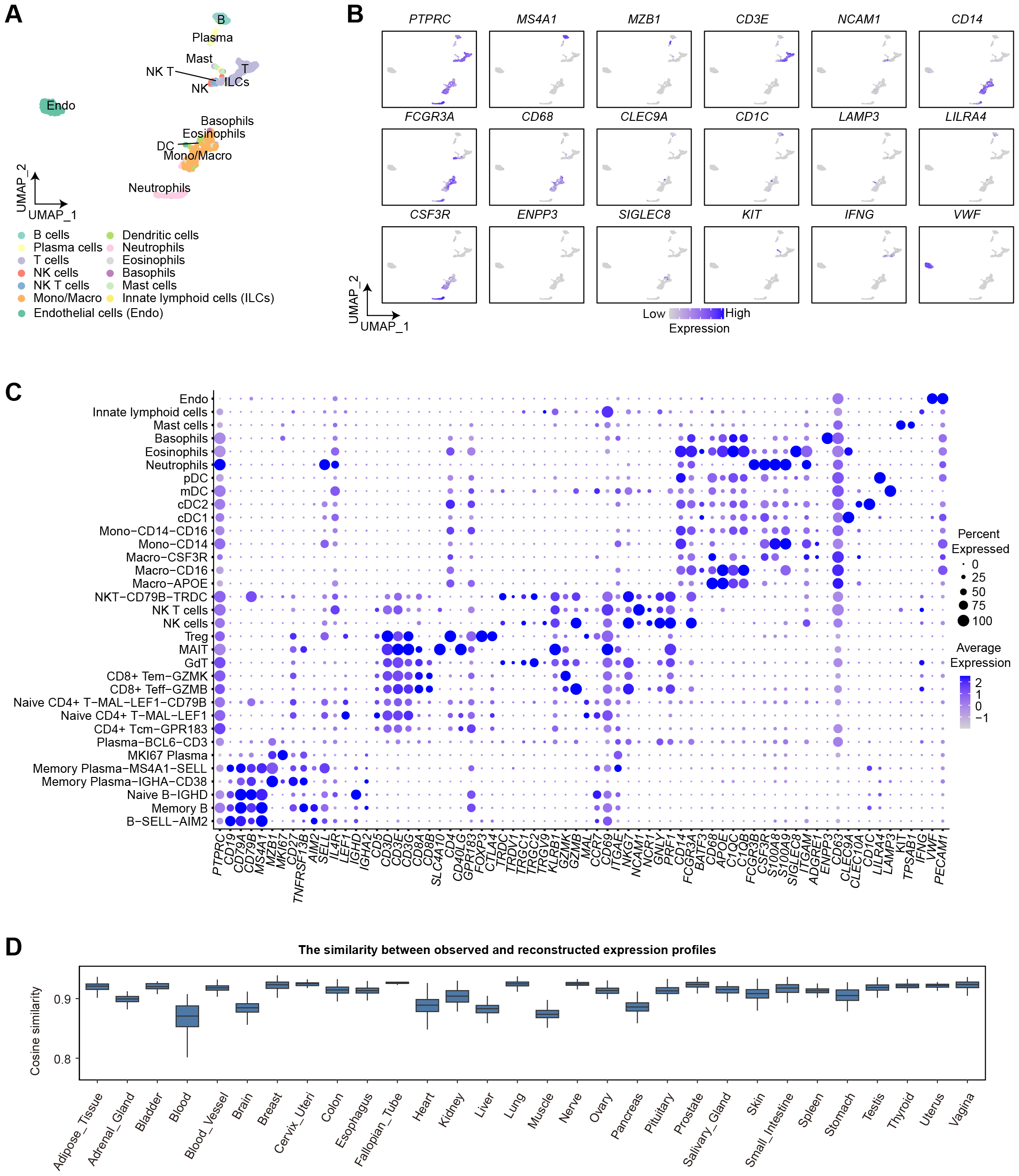
Annotation of scRNA-seq data and accuracy of deconvolution. A, UMAP visualization of 1247 scRNA-seq profiles colored by major cell types. B, Expression of signature genes of 13 major cell types. C, Dot plot indicating the expression of signature genes of 33 cell clusters. D, Cosine similarities of tissue samples between observed and reconstructed expression profiles in 30 tissues.

### Validations based on literature data

Sender et al conducted a literature survey of cellular densities (cells/g) of the primary immune cell types in various tissues based on histology or flow cytometry techniques ^11^. Here, we validated the high accuracy of RedeImmunoMap by correlating our estimated abundance with the corresponding cellular densities, ensuring the reliability of our findings (**Fig. 3**). The pearson correlation between the predicted immune abundance and immune cell density reached 0.93 (*P*-value=4.1e-05) in 11 tissues, proving the high precision of RedeImmunoMap (**Fig. 3A**). In views of the specificalities of blood and adipose tissue on measurement of cell densities, we didn’t consider them here. Furthermore, we performed pearson correlation analysis on various immune cell types including B cells, T cells, monocytes/macrophages, neutrophils, eosinophils and mast cells (**Fig. 3B-C**). For the lymphocytes, the correlations reached 0.72 (*P*-value=0.012) and 0.77 (*P*-value=0.0056) respectively for B and T cells (**Fig. 3B)**. As for the myeloid cells, we also observed high correlations between the predicted abundance and corresponding cell density with pearson correlations of 0.83 (*P*-value =0.0014) for monocytes/macrophages, 0.78 (*P*-value =0.021) for neutrophils, 0.9 (*P*-value =0.038) and 0.92 (*P*-value=0.0035) for mast cells (**Fig. 3C)**. Overall, we successfully reconstructed immune cellular compartments of human tissues (**Fig. 3D**), holding the potential to yield profound insights into human immunology and health.

**Fig. 3.**
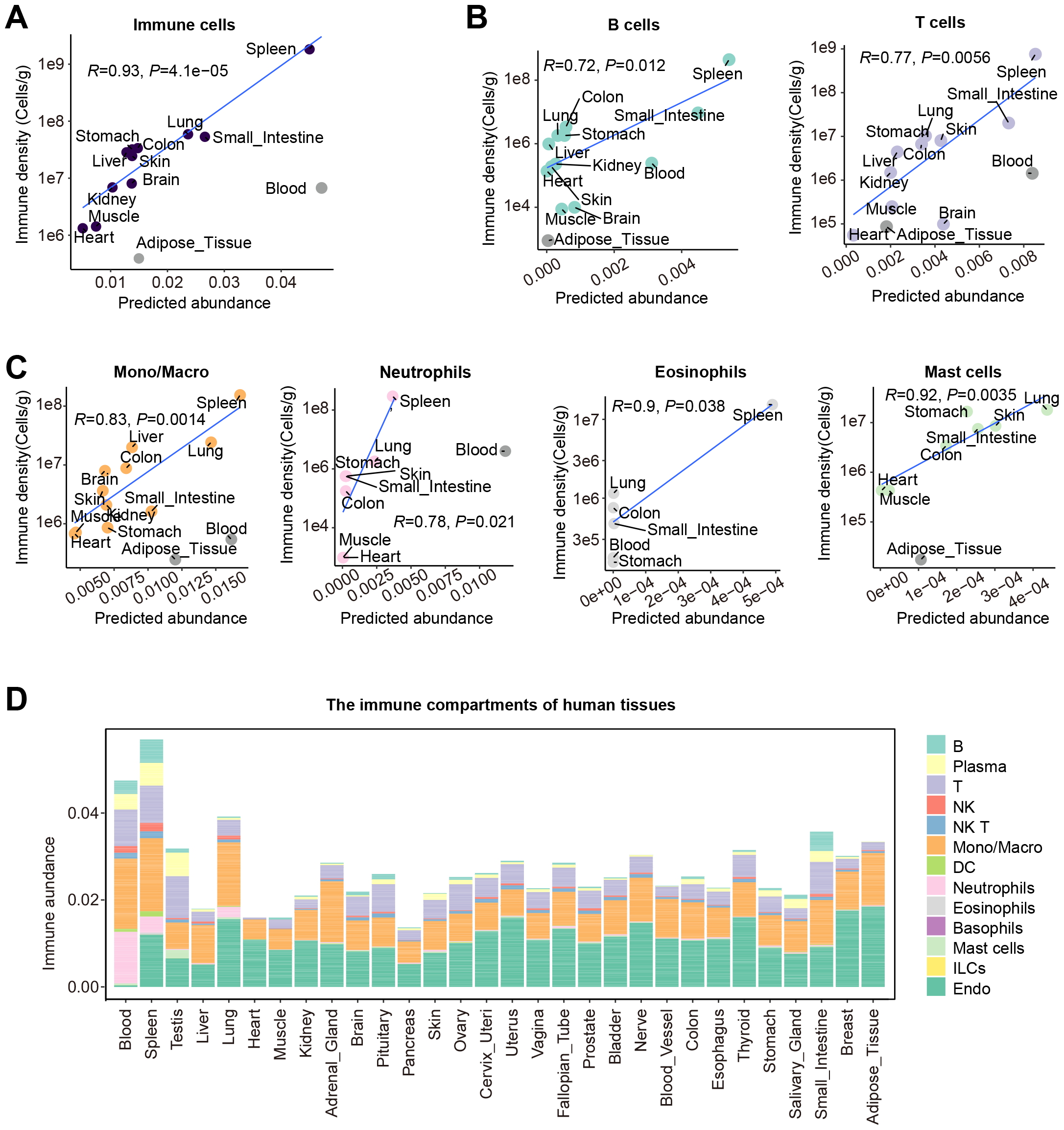
Validation of estimated immune abundance. A, Pearson correlation between estimated immune abundance and corresponding immune cell densities published by Sender et al. B-C, Pearson correlation between estimated immune subtype abundance and corresponding cell densities published by Sender et al. D, Bar plot showing the cellular compartments of each tissue. The abundance of each cell state was averaged for each tissue.

### Distribution of endothelial cells in human tissues across different sexes and ages

Endothelial cells form the barrier between blood vessels and tissue as well as engage in intimate crosstalk with immune cells, thereby exerting a crucial role in regulating immune trafficking, stimulating immune responses, and inhibiting immune activation ^5,12^. Therefore, a thorough knowledge of the distribution of endothelial cells across human tissues is likely to contribute to our enhanced comprehension of immune functions. We compiled the abundance of endothelial cells according to their ranks, showing high heterogeneity across different tissues (**Fig. 4**). We observed adipose tissue and breast had the highest abundance of endothelial cells, followed by thyroid, uterus and lung. Comparatively, blood rarely harbors endothelial cells while liver, pancreas and testis exhibit a notably lower abundance of endothelial cells than other tissues (**Fig. 4A**). Furthermore, we investigated the distribution of endothelial cells in the males and females across diverse tissues (**Fig. 4B**). We noticed that endothelial cell abundance had no significant difference between the males and females in the majority of tissues. Notably, a significantly higher abundance of endothelial cells were observed in adipose tissue, breast, and thyroid among the males compared to the females, whereas brain exhibits a lower endothelial cell abundance in the males. Additionally, we characterized the distribution of endothelial cells in different age groups across tissues (**Fig. 4C**). A strong association of endothelial cell abundance with age was observed in breast, lung, nerve, cervix uteri, colon, ovary, muscle, brain, skin and liver by ANOVA analysis, revealing that these tissues are susceptible to endothelial senescence, which can potentially lead to cardiovascular disease as a result of progressive endothelial dysfunction ^13^.

**Fig. 4.**
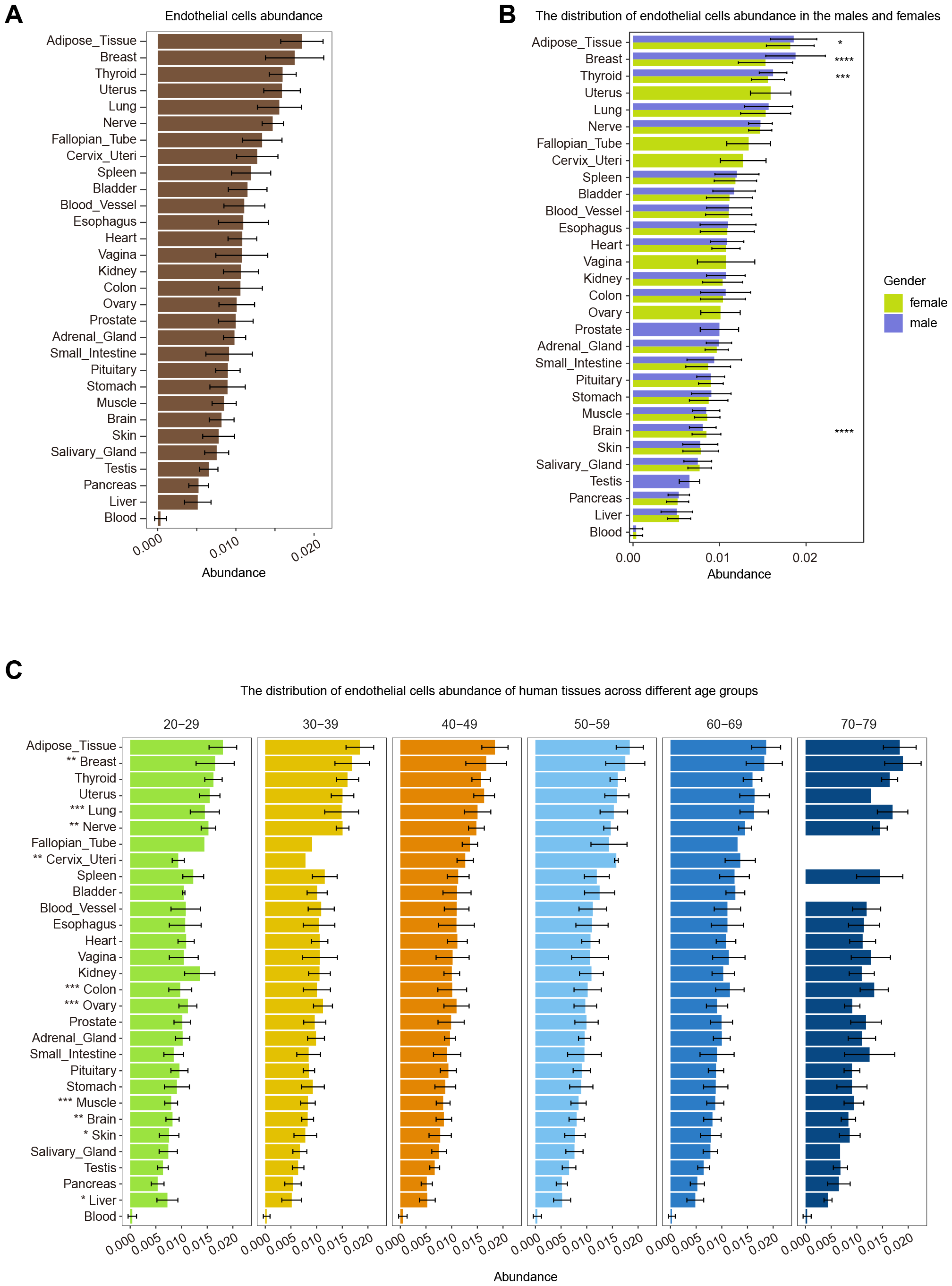
Distribution of endothelial cell abundance across different tissues, sexes and ages. A, Bar plot showing the distribution of endothelial cell abundance of 30 tissues ranked by their averaged endothelial cell abundance. Error bars were plotted by mean±SD for A, B and C. The endothelial cell abundance of each RNA-seq sample was aggregated for all endothelial cell states for A, B and C. B, Comparisons of endothelial cell abundance between the males and females (Wilcoxon rank-sum test). C, Comparisons of endothelial cell abundance across different age groups (ANOVA test). The significant level was correspondingly marked in the front of tissue names. \**P* < 0.05, \*\**P* < 0.01, \*\*\**P* < 0.001, \*\*\*\**P* < 0.0001.

### Immune cell abundance of human body across different sexes and ages

Immune cells recognize and combat harmful substances, pathogens, and cellular changes throughout the tissues of our bodies, protecting humans from various pathologies ^14^. Previous studies have reported that age and gender function as the most powerful drivers of immune variation ^15^. However, how immune cells distribute over our bodies across different sexes and ages has not been well studied before, though previous studies have investigated partial tissues ^16^. Here, we provided the most comprehensive distribution of immune cells over 30 human tissues (**Fig. 5**). As presented in Fig. 5A, blood, spleen, small intestine are the most immunogenic tissues with remarkably abundant immune cells, as widely reported in previous studies ^11,17^ (**Fig. 5A**). Notably, we observed a high abundance of immune cells in testis as an immune privileged organ only second to small intestine, which has not been reported yet ^18^. Comparatively, heart exhibit a lowest abundance of immune cells, as reported in the previous study that immune cells occupied the smallest proportion of the whole in heart ^19^. We further compared the immune cell abundance between the males and females, showing tissue-specific significant differences (**Fig. 5B**). The women tend to have a higher abundance of immune cells in thyroid, adipose tissue, breast, kidney, muscle and heart while the men exhibit a greater concentration of immune cells in salivary gland and pancreas. ANOVA analysis revealed a significant correlation of immune cell abundance with age in 40% tissues (**Fig. 5C**). We observed an age-dependent decrease in immune cell abundance in blood, small intestine, stomach and liver, contrasted by an age-related increase in nerve, prostate and blood vessel. Briefly, we presented a comprehensive overview of immune cell abundance in human tissues and associations with sex and age.

**Fig. 5.**
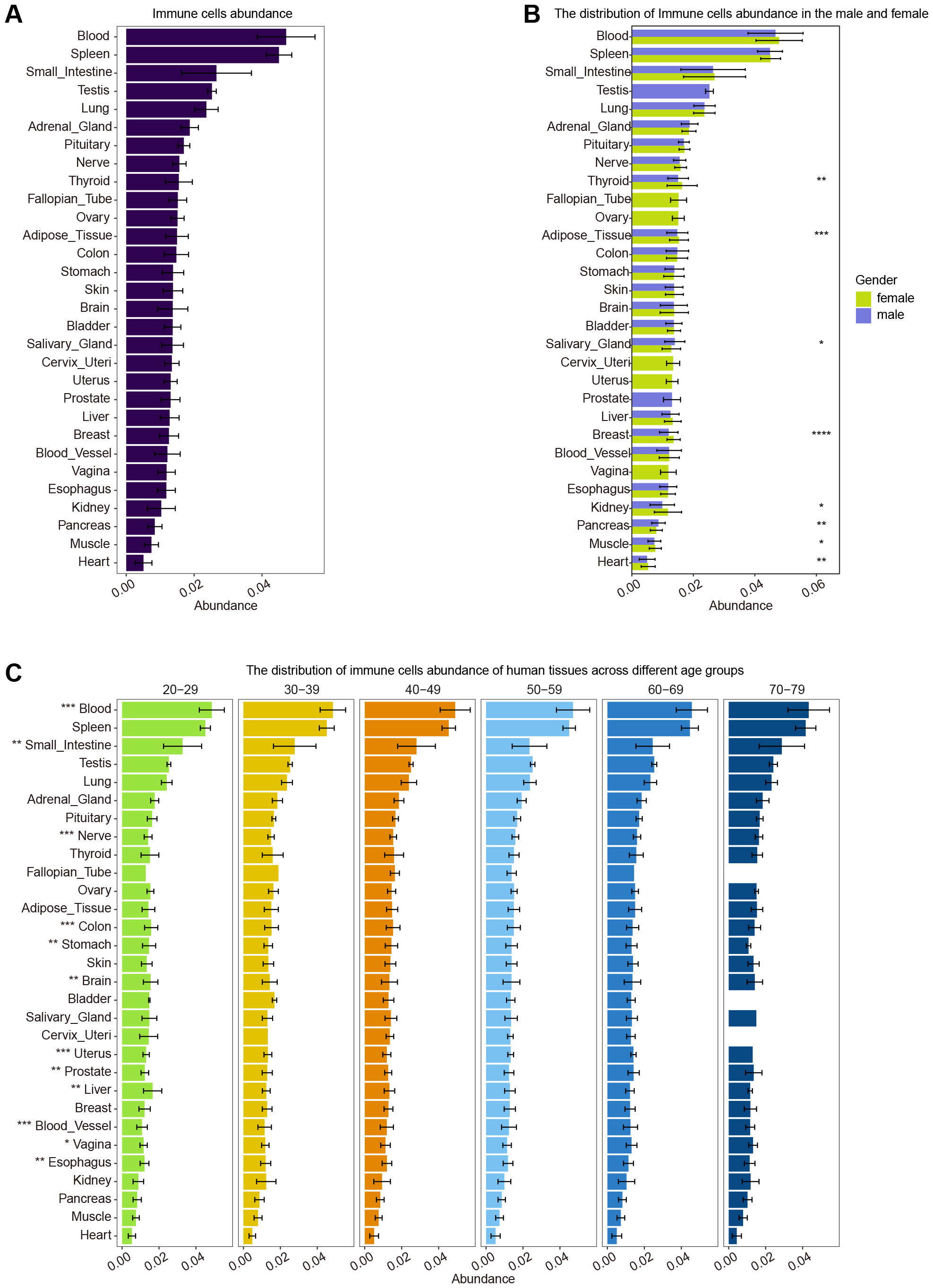
Distribution of immune cell abundance across different tissues, sexes and ages. A, Bar plot showing the distribution of immune cell abundance of 30 tissues ranked by their averaged immune cell abundance. Error bars were plotted by mean±SD for A, B and C. The immune cell abundance of each RNA-seq sample was aggregated for all immune cell states for A, B and C. B, Comparisons of immune cell abundance between the males and females (Wilcoxon rank-sum test). C, Comparisons of immune cell abundance across different age groups (ANOVA test). The significant level was correspondingly marked in the front of tissue names. \**P* < 0.05, \*\**P* < 0.01, \*\*\**P* < 0.001, \*\*\*\**P* < 0.0001.

### Abundance of different immune cell types across different tissues, sexes and ages

Furthermore, we explored the abundance of 12 immune cell subtypes across 30 tissues including B cells, plasma cells, T cells, NK cells, NKT cells, mono/macro, DC, neutrophils, eosinophils, basophils, mast cells and ILCs (**Fig. S1-S12**).

We observed a significant enrichment of B cells, plasma cells, T cells in blood, spleen, small intestine and testis but depletion in heart (**Fig. S1-S5**). B cells also exhibit high abundance and significant correlation with sex in pituitary and salivary gland as well as significantly correlate with age in about 33% tissues including small intestine, brain, colon, prostate, uterus, vagina and so on (**Fig. S1**). Plasma cells show significant sex-related difference in salivary gland and breast as well as have a strong association with age in salivary gland, small intestine, colon, cervix uteri, brain and breast (**Fig. S2**). T cells play a substantial role in pituitary, ovary and thyroid, interplaying with sex in brain, skin and breast as well as with age in 40% tissues including small intestine, uterus, brain, skin and nerve (**Fig. S3**). spleen and blood besides lung, small intestine and prostate harbor abundant NK cells. We observed significant difference of NK cell abundance across different tissues, sexes and ages in spleen, adipose tissue, skin and esophagus (**Fig. S4**). NKT cells exhibit a significant enrichment in spleen, blood, uterus, pituitary and cervix uteri. Sex-related difference of NKT cell abundance was observed in pituitary, nerve, thyroid, breast and muscle while age-related difference was significantly enriched in blood, pituitary, nerve, brain, blood vessel, adipose tissue and muscle (**Fig. S5**). Myeloid cells play major roles in innate immunity by secretion of inflammatory cytokines and chemokines upon pathogen invasion ^4^. Hence, we characterized the cell type abundance of myeloid subsets across different tissues (**Fig. S6-S11**). Mono/macro as the largest myeloid populations are preferentially enriched in spleen, blood, lung, adrenal gland, adipose tissue and small intestine. Significant difference of mono/macro abundance was observed in adipose tissue, thyroid, skin and heart across different sexes while blood, lung, small intestine, blood vessel, liver, brain, skin and ovary exhibit age-related difference of mono/macro abundance (**Fig. S6**). Dendritic cells abundance displays a dominantly enrichment in spleen, blood, lung and salivary gland and age-related difference in blood, lung and breast (**Fig. S7**). Neutrophils, basophils, eosinophils were mainly detected in a few tissues including blood, spleen or lung (**Fig. S8-S10**). We found a large amount of mast cells resided in testis far more than any other tissue, followed by lung, blood, skin, bladder and small intestine. The men have a significantly higher abundance of mast cells than women in lung, small intestine, esophagus, stomach and blood vessel. Significant age-related difference of mast cell abundance was observed in lung, blood, prostate, colon, brain, nerve, ovary and spleen (**Fig. S11**). Notably, we found numerous ILCs in ovary and more than 40% tissues exhibit significant gender-related difference across different sexes and 30% tissues display significant age-related difference across different ages (**Fig. S12**).

In conclusion, spleen, blood, small intestine and testis are important organs with strong adaptive or initiate immunities while heart and muscle are immunologically isolated organs. Lung is a specific immune organ that harbor abundant NK cells and myeloid cells including dendritic cells, neutrophils, basophils and mast cells. Ovary and pituitary gather plenty of lymphoid cells including B cells, plasma cells and T cells or ILCs. Our results serve as a foundation for further exploration into the tissue-specific, sex-specific and age-specific immunities of human body at a global scale.

### Immune similarities among human tissues

The immune system operates as an intricate and integrated network composed of various tissues, corporately performing tissue-specific functions of immune regulation and responses ^20^. However, immune associations between different human tissues have not been scientifically investigated before. Here, we introduced the similarities and differences between 30 human tissues based on the abundance of immune plus endothelial cell states (**Fig. 6**). We first visualized the immune associations between different tissues in two dimensions using t-distributed stochastic neighbor embedding (t-SNE) (**Fig. 6A**) and simultaneously compared the distribution of cell abundance across various tissues for each individual cell type (**Fig. 6B**). Furthermore, we comprehensively depicted the pearson correlation of immune plus endothelial cell states between different tissues (**Fig. 6C**). Accordingly, we concluded 14 different tissue modules according to their immune similarities or biological functions. Blood, spleen and testis exhibit extremely specific immunities with unique immune compartments, reflected in high abundance of different immune subtypes. Liver and lung are also special organs showing faint similarities with other tissues. Notably, we found that the tissues with relevant biological functions or similar materials tend to have similar immune compartments. For examples, the correlation reaches 0.82 between brain and pituitary, showing a minor enrichment of lymphoid cells. The tissues of the reproductive system obtain an averaged correlation of 0.9 around, with a slight enrichment in T and NK cells. Likewise, the digestive system demonstrates notable similarities in immune compartments, containing more B and plasma cells. Breast and adipose tissue display strikingly similar immune properties and high abundance of endothelial cells. Overall, our results comprehensively elucidate the immune associations across various human tissues, thereby enhancing our holistic understanding of the intricate mechanisms underlying the human immune system.

**Fig. 6.**
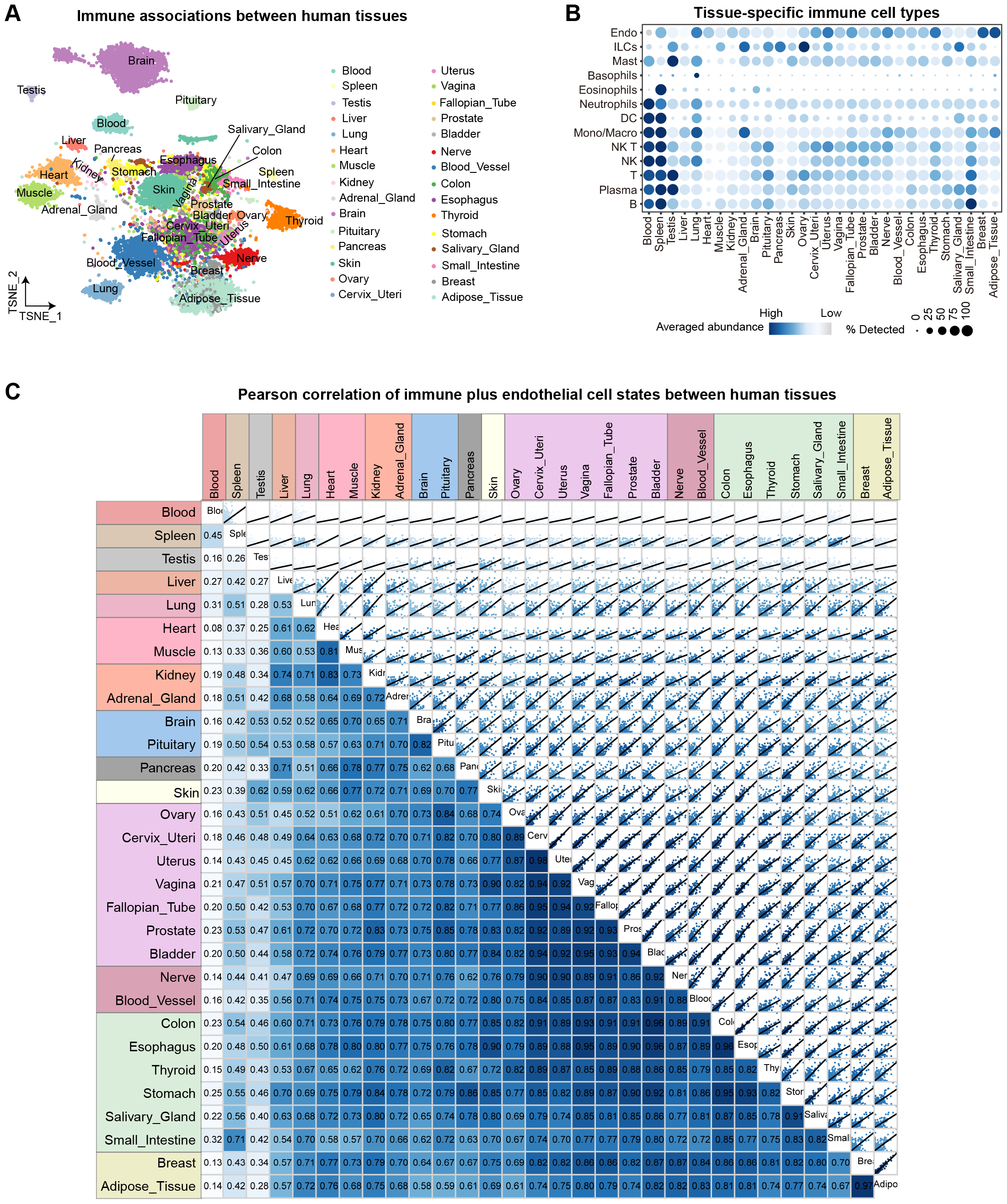
Immune associations between human tissues. A, t-SNE visualization of 17382 RNA-seq samples based on their cell state abundance. B, Dot plot showing tissue-specific abundant cell types colored by scaled mean abundance across tissues for each major cell type. The size of points indicted the percentage of samples detecting the major cell type. C. Pearson correlation between different tissues plotted by corrgram. The upper panels indicated the scatter plots of averaged cell abundance for 1247 cell states between two tissues. The lower panels indicated the pearson correlation values of cell state abudnance between two tissues. The color of each square panel indicated corresponding pearson correlation values.

## Discussion

The integration and subsequent analysis of scRNA-seq and bulk RNA-seq datasets serve as an asset in elucidating the comprehensive map of cellular landscape and intricate interactomes within the tissues of our scientific interest. Here, we provide a novel sight about how to depict the intricate immune system by existing methods and fully utilize valuable and large-scale bulk RNA-seq datasets in our study. We effectively demonstrated the accuracy and reliability of RedeImmunoMap by validating on published datasets based on histology or flow cytometry techniques. Then, we thoroughly displayed the distribution of immune and endothelial abundance as well as immune subtype abundance across 30 tissues. We found that blood, spleen, small intestine and testis were the major immune organs, harboring the largest abundance of immune cells. For specific immune cell types/subtypes, the highly-enriched organs/tissues alter accordingly, suggesting regional immune specificities. We classified 30 tissues into 14 different tissue modules according to their immune characteristics, revealing that blood, spleen, testis, liver, lung, and some other organs/tissues are uniquely-defined immune compartments, while the reproductive system and digestive system that are characterized by tube-or luminal morphology demonstrate notable immune similarities. Overall, RedeImmunoMap comprehensively elucidates the immune compositions across various human tissues and their associations with sex and age, which provides a rich data resource for investigating human immune system from a holistic perspective.

## Methods

### Public datasets

The bulk RNA-seq expression profiles and corresponding metadata information were downloaded from https://www.gtexportal.org/home/datasets, including 17382 samples from 30 tissues. The scRNA-seq data originated from the Tabula Sapiens were downloaded from https://figshare.com/articles/dataset/Tabula_Sapiens_release_1_0/14267219, including 483,152 cells from 24 human tissues and organs. The measured immune cell densities based on histology or flow cytometry techniques were collected from Sender et al ^11^.

### Deconvolution of RNA-seq data

The bulk RNA-seq expression values were estimated by the Transcripts Per Million (TPM) measure, then log_2_ transformed. To simplify the computational burden, 5002 cell states were sampled from 483,152 cells via Redeconve based on their original cell type annotations. Further, the log-transformed bulk RNA-seq expression matrix and sampled scRNA-seq expression matrix were together inputted into Redeconve to estimate the abundance of each cell state in each RNA-seq sample. 19965 highly variable genes were saved by Redeconve using the function of *gene*.*filter*. The obtained abundance matrix was used for subsequent analysis.

### scRNA-seq data annotation

The sampled scRNA-seq data containing 5002 single cells were analyzed by Seurat v5 workflow and integrated by Harmony ^21,22^. Data normalization, preprocessing, and clustering were performed. Immune and endothelial clusters were annotated and selected via the corresponding signature genes including *PTPRC, VWF, PECAM1*. The cells expressing *PTPRC* were also annotated immune cells. 2920 immune plus endothelial cells were picked and clustered into 15 cell clusters. We further annotated immune clusters into B cells (*MS4A1*), plasma cells (*MZB1*), T cells (*CD3*), NK cells (*NCAM1*), NKT cells (*CD3*+*NCAM1*+), monocytes/macrophages (*CD14*/*CD16*), dendritic cells (*CLEC9A*/*CD1C*/*LILRA4*/*LAMP3*), neutrophils (*FCGR3B*/*CSF3R*), eosinophils (*SIGLEC8*), basophils (*ENPP3*) and mast cells (*KIT*). Initiate lymphoid cells were annotated based on original paper. Furthermore, we clustered B cells, plasma cells, T cells and monocytes/macrophages respectively. Finally, we picked detected cell states by Redeconve and totally obtained 999 immune and 248 endothelial cell states clustered into 13 major cell types and 33 cell subtypes.

### Immune similarity analysis across tissues

The RNA-seq samples were visualized in two dimensions using t-distributed stochastic neighbor embedding (t-SNE) based on their abundance of 1247 cell states. Furthermore, the abundance of immune and endothelial cell states in each tissue was averaged and then used for pearson correlation analysis. We visualized the correlations between 30 tissues via R package “corrgram”^23^.

## Acknowledgements

This work was supported by the National Natural Science Foundation of China (32022016 to X.R., 92159305 to X.R., and 31991171 to X.R.), National Key R&D Program of China (2022YFC3400904 to X.R.), and Changping Laboratory.

## Author contributions

X.R conceives and supervises this project. R.G. conducts the deconvolution and subsequent analysis of different tissues regarding the immune composition and associations with sex and age with the contribution from X.J. R.G. and X.R. wrote the manuscript together.

## Competing interests

The authors declare no competing interests.

**Supplementary Fig. 1. Distribution of B cell abundance across different tissues, sexes and ages**. A, Bar plot showing the distribution of B cell abundance of 30 tissues ranked by their averaged B cell abundance. Error bars were plotted by mean±SD for A, B and C. The B cell abundance of each RNA-seq sample was aggregated for all B cell states for A, B and C. B, Comparisons of B cell abundance between the males and females (Wilcoxon rank-sum test). C, Comparisons of B cell abundance across different age groups (ANOVA test). The significant level was correspondingly marked in the front of tissue names. \**P* < 0.05, \*\**P* < 0.01, \*\*\**P* < 0.001, \*\*\*\**P* < 0.0001.

**Supplementary Fig. 2. Distribution of plasma cell abundance across different tissues, sexes and ages**. A, Bar plot showing the distribution of plasma cell abundance of 30 tissues ranked by their averaged plasma cell abundance. Error bars were plotted by mean±SD for A, B and C. The plasma cell abundance of each RNA-seq sample was aggregated for all plasma cell states for A, B and C. B, Comparisons of plasma cell abundance between the males and females (Wilcoxon rank-sum test). C, Comparisons of plasma cell abundance across different age groups (ANOVA test). The significant level was correspondingly marked in the front of tissue names. \**P* < 0.05, \*\**P* < 0.01, \*\*\**P* < 0.001, \*\*\*\**P* < 0.0001.

**Supplementary Fig. 3. Distribution of T cell abundance across different tissues, sexes and ages**. A, Bar plot showing the distribution of T cell abundance of 30 tissues ranked by their averaged T cell abundance. Error bars were plotted by mean±SD for A, B and C. The T cell abundance of each RNA-seq sample was aggregated for all T cell states for A, B and C. B, Comparisons of T cell abundance between the males and females (Wilcoxon rank-sum test). C, Comparisons of T cell abundance across different age groups (ANOVA test). The significant level was correspondingly marked in the front of tissue names. \**P* < 0.05, \*\**P* < 0.01, \*\*\**P* < 0.001, \*\*\*\**P* < 0.0001.

**Supplementary Fig. 4. Distribution of NK cell abundance across different tissues, sexes and ages**. A, Bar plot showing the distribution of NK cell abundance of 30 tissues ranked by their averaged NK cell abundance. Error bars were plotted by mean±SD for A, B and C. The NK cell abundance of each RNA-seq sample was aggregated for all NK cell states for A, B and C. B, Comparisons of NK cell abundance between the males and females (Wilcoxon rank-sum test). C, Comparisons of NK cell abundance across different age groups (ANOVA test). The significant level was correspondingly marked in the front of tissue names. \**P* < 0.05, \*\**P* < 0.01, \*\*\**P* < 0.001, \*\*\*\**P* < 0.0001.

**Supplementary Fig. 5. Distribution of NKT cell abundance across different tissues, sexes and ages**. A, Bar plot showing the distribution of NKT cell abundance of 30 tissues ranked by their averaged NKT cell abundance. Error bars were plotted by mean±SD for A, B and C. The NKT cell abundance of each RNA-seq sample was aggregated for all NKT cell states for A, B and C. B, Comparisons of NKT cell abundance between the males and females (Wilcoxon rank-sum test). C, Comparisons of NKT cell abundance across different age groups (ANOVA test). The significant level was correspondingly marked in the front of tissue names. \**P* < 0.05, \*\**P* < 0.01, \*\*\**P* < 0.001, \*\*\*\**P* < 0.0001.

**Supplementary Fig. 6. Distribution of monocytes/macrophages abundance across different tissues, sexes and ages**. A, Bar plot showing the distribution of monocytes/macrophages cell abundance of 30 tissues ranked by their averaged monocytes/macrophages cell abundance. Error bars were plotted by mean±SD for A, B and C. The monocytes/macrophages cell abundance of each RNA-seq sample was aggregated for all monocytes/macrophages cell states for A, B and C. B, Comparisons of monocytes/macrophages cell abundance between the males and females (Wilcoxon rank-sum test). C, Comparisons of monocytes/macrophages cell abundance across different age groups (ANOVA test). The significant level was correspondingly marked in the front of tissue names. \**P* < 0.05, \*\**P* < 0.01, \*\*\**P* < 0.001, \*\*\*\**P* < 0.0001.

**Supplementary Fig. 7. Distribution of dendritic cell abundance across different tissues, sexes and ages**. A, Bar plot showing the distribution of dendritic cell abundance of 30 tissues ranked by their averaged dendritic cell abundance. Error bars were plotted by mean±SD for A, B and C. The dendritic cell abundance of each RNA-seq sample was aggregated for all dendritic cell states for A, B and C. B, Comparisons of dendritic cell abundance between the males and females (Wilcoxon rank-sum test). C, Comparisons of dendritic cell abundance across different age groups (ANOVA test). The significant level was correspondingly marked in the front of tissue names. \**P* < 0.05, \*\**P* < 0.01, \*\*\**P* < 0.001, \*\*\*\**P* < 0.0001.

**Supplementary Fig. 8. Distribution of neutrophils abundance across different tissues, sexes and ages**. A, Bar plot showing the distribution of neutrophils abundance of 30 tissues ranked by their averaged neutrophils abundance. Error bars were plotted by mean±SD for A, B and C. The neutrophils abundance of each RNA-seq sample was aggregated for all neutrophils states for A, B and C. B, Comparisons of neutrophils abundance between the males and females (Wilcoxon rank-sum test). C, Comparisons of neutrophils abundance across different age groups (ANOVA test). The significant level was correspondingly marked in the front of tissue names. \**P* < 0.05, \*\**P* < 0.01, \*\*\**P* < 0.001, \*\*\*\**P* < 0.0001.

**Supplementary Fig. 9. Distribution of eosinophils abundance across different tissues, sexes and ages**. A, Bar plot showing the distribution of eosinophils abundance of 30 tissues ranked by their averaged eosinophils abundance. Error bars were plotted by mean±SD for A, B and C. The eosinophils abundance of each RNA-seq sample was aggregated for all eosinophils states for A, B and C. B, Comparisons of eosinophils abundance between the males and females (Wilcoxon rank-sum test). C, Comparisons of eosinophils abundance across different age groups (ANOVA test). The significant level was correspondingly marked in the front of tissue names. \**P* < 0.05, \*\**P* < 0.01, \*\*\**P* < 0.001, \*\*\*\**P* < 0.0001.

**Supplementary Fig. 10. Distribution of basophils abundance across different tissues, sexes and ages**. A, Bar plot showing the distribution of basophils abundance of 30 tissues ranked by their averaged basophils abundance. Error bars were plotted by mean±SD for A, B and C. The basophils abundance of each RNA-seq sample was aggregated for all basophils states for A, B and C. B, Comparisons of basophils abundance between the males and females (Wilcoxon rank-sum test). C, Comparisons of basophils abundance across different age groups (ANOVA test). The significant level was correspondingly marked in the front of tissue names. \**P* < 0.05, \*\**P* < 0.01, \*\*\**P* < 0.001, \*\*\*\**P* < 0.0001.

**Supplementary Fig. 11. Distribution of mast cell abundance across different tissues, sexes and ages**. A, Bar plot showing the distribution of mast cell abundance of 30 tissues ranked by their averaged mast cell abundance. Error bars were plotted by mean±SD for A, B and C. The mast cell abundance of each RNA-seq sample was aggregated for all mast cell states for A, B and C. B, Comparisons of mast cell abundance between the males and females (Wilcoxon rank-sum test). C, Comparisons of mast cell abundance across different age groups (ANOVA test). The significant level was correspondingly marked in the front of tissue names. \**P* < 0.05, \*\**P* < 0.01, \*\*\**P* < 0.001, \*\*\*\**P* < 0.0001.

**Supplementary Fig. 12. Distribution of initiate lymphoid cell abundance across different tissues, sexes and ages**. A, Bar plot showing the distribution of initiate lymphoid cell abundance of 30 tissues ranked by their averaged initiate lymphoid cell abundance. Error bars were plotted by mean±SD for A, B and C. The initiate lymphoid cell abundance of each RNA-seq sample was aggregated for all initiate lymphoid cell states for A, B and C. B, Comparisons of initiate lymphoid cell abundance between the males and females (Wilcoxon rank-sum test). C, Comparisons of initiate lymphoid cell abundance across different age groups (ANOVA test). The significant level was correspondingly marked in the front of tissue names. \**P* < 0.05, \*\**P* < 0.01, \*\*\**P* < 0.001, \*\*\*\**P* < 0.0001.

